# *Listeria monocytogenes* GlmR is an accessory uridyltransferase essential for cytosolic survival and virulence

**DOI:** 10.1101/2021.10.27.466214

**Authors:** Daniel A. Pensinger, Kimberly V. Gutierrez, Hans B. Smith, William J.B. Vincent, David S. Stevenson, Katherine A. Black, Krizia M. Perez-Medina, Joseph P. Dillard, Kyu Y. Rhee, Daniel Amador-Noguez, TuAnh N Huynh, John-Demian Sauer

**Author notes:** equal contributions (Daniel A. Pensinger, Kimberly V. Gutierrez). Corresponding Author: Dr. John-Demian Sauer, Department of Medical Microbiology and Immunology, University of Wisconsin-Madison 1550 Linden Dr. Rm 4203, Madison WI, 53706 USA.

## Abstract

The cytosol of eukaryotic host cells is an intrinsically hostile environment for bacteria. Understanding how cytosolic pathogens adapt to and survive in the cytosol is critical to developing novel therapeutic interventions for these pathogens. The cytosolic pathogen *Listeria monocytogenes* requires *glmR* (previously known as *yvcK*), a gene of unknown function, for resistance to cell wall stress, cytosolic survival, inflammasome avoidance and ultimately virulence *in vivo*. A genetic suppressor screen revealed that blocking utilization of UDP-GlcNAc by a non-essential wall teichoic acid decoration pathway restored resistance to cell wall stress and partially restored virulence of Δ*glmR* mutants. In parallel, metabolomics revealed that Δ*glmR* mutants are impaired in the production of UDP-GlcNAc, an essential peptidoglycan and wall teichoic acid (WTA) precursor. We next demonstrated that purified GlmR can directly catalyze the synthesis of UDP-GlcNAc from GlcNAc-1P and UTP, suggesting that it is an accessory uridyltransferase. Biochemical analysis of GlmR orthologues suggest that uridyltransferase activity is conserved. Finally, mutational analysis resulting in a GlmR mutant with impaired catalytic activity demonstrated that uridyltransferase activity was essential to facilitate cell wall stress responses and virulence in vivo. Taken together these studies indicate that GlmR is an evolutionary conserved accessory uridyltransferase required for cytosolic survival and virulence of *L. monocytogenes*.

**Importance:** Bacterial pathogens must adapt to their host environment in order to cause disease. The cytosolic bacterial pathogen *Listeria monocytogenes* requires a highly conserved protein of unknown function, GlmR (previously known as YvcK) to survive in the host cytosol. GlmR is important for resistance to some cell wall stresses and is essential for virulence. The Δ*glmR* mutant is deficient in production of an essential cell wall metabolite, UDP-GlcNAc, and suppressors which increase metabolite levels also restore virulence. Purified GlmR can directly catalyze the synthesis of UDP-GlcNAc and this enzymatic activity is conserved in pathogens from Firmicutes and Actinobacteria phyla. These results highlight the importance accessory cell wall metabolism enzymes in responding to cell wall stress in a variety of bacterial pathogens.

## Introduction

Bacterial pathogens encounter a variety of stresses throughout the course of infection ranging from nutritional stresses, redox stresses and cell wall stresses. Specifically, the mammalian cytosol restricts the survival and replication of bacteria that are not adapted for that niche (1–7). To protect the cytosol, the host utilizes a variety of known and unknown cell autonomous defenses (CADs) that directly target bacterial survival (8, 9). Despite this, canonical cytosolic pathogens such as *Listeria monocytogenes* can replicate efficiently in this environment. These bacterial pathogens have developed adaptions to survive host imposed stresses in the cytosol (10), acquire necessary nutrients (11), and divert or subvert innate immune defenses (12, 13). Although many of the adaptations that allow cytosol adapted pathogens to endure host defenses and stress in the cytosol remain unknown, recent genetic screens have identified some bacterial genes that contribute to cytosolic survival, however the molecular function of many of these genes remains unknown (7, 14, 15).

A number of virulence factors essential for cytosolic survival of *L. monocytogenes*, a highly cytosol adapted pathogen, have recently been identified (4, 14, 16, 17). One such protein, GlmR (also known as YvcK or CuvA), is a highly conserved protein found in firmicutes and actinobacteria. GlmR and its homologues are dispensable for growth in nutrient rich media, but are essential for growth on gluconeogenic carbon sources and in the presence of cell wall stress (16, 18, 19). Consistent with these functions, *L. monocytogenes* GlmR expression is also induced in response to cell wall stress (16). Finally, *L. monocytogenes* GlmR is required for cytosolic survival and replication in host cells (14), and is required for virulence of both *L. monocytogenes* and *Mycobacterium tuberculosis in vivo* (16, 19, 20). In *S. aureus* GlmR is predicted to be essential, even in rich media in the absence of cell wall stress (21). Despite the striking phenotypes of Δ*glmR* mutants in a variety of organisms, molecular function(s) of the protein remain largely unknown.

How GlmR contributes to cell wall stress responses and virulence remains largely unknown, however, GlmR was recently described to bind to the essential cell wall precursor UDP-N-acetyl-glucosamine (UDP-GlcNAc) (22). UDP-GlcNAc is required for the synthesis of peptidoglycan, wall teichoic acid in Firmicutes, and arbinogalactan in *M. tuberculosis* (23–25). In *B. subtilis*, GlmR was found to interact with GlmS, one of three highly conserved proteins necessary for UDP-GlcNAc synthesis (26). To characterize the function of GlmR in *L. monocytogenes* we first utilized a genetic suppressor screen to identify second site mutations that restored lysozyme resistance of the Δ*glmR* mutant. Two independent suppressor mutants A increased pools of UDP-GlcNAc ultimately restoring cell wall stress responses and virulence of Δ*glmR* mutants. In parallel, untargeted metabolomics revealed that Δ*glmR* mutants are deficient in UDP-GlcNAc. We were unable to detect interactions between *L. monocytogenes* GlmR and its cognate GlmS as previously reported in *B. subtilis* and instead found that purified GlmR, and its orthologues, demonstrate uridyltransferase activity that can catalyze the synthesis UDP-GlcNAc from UTP and GlcNAc-1P. Finally, mutational analysis demonstrated that GlmR uridyltransferase activity is necessary to promote cell wall stress responses and virulence *in vivo*. Together our data suggests that GlmR is an accessory uridyltransferase that is upregulated to deal with cell wall stress such as that encountered by *L. monocytogenes* during cytosolic replication.

## Results

### Inhibition of non-essential decoration of wall teichoic acid with GlcNAc rescues cell wall stress defects of the Δ*glmR* mutant

*L. monocytogenes* GlmR is essential for cytosolic survival and virulence, is upregulated in the context of lysozyme stress and is necessary for resistance to lysozyme (16). To understand how GlmR contributes to cell wall stress responses and virulence we performed a lysozyme resistance suppressor selection using a Himar1 mariner-based transposon mutant library in a Δ*glmR* mutant background. Twenty unique transposon insertions across fifteen unique genes suppressed the Δ*glmR* mutant’s lysozyme sensitivity (Table 1). The suppressors represent a diverse set of cellular processes that likely contribute to lysozyme resistance in a variety of ways including mechanisms that are both generic and GlmR specific. Mutations which generically upregulate stress response pathways may not be useful for understanding GlmR function. Therefore, to prioritize cell wall stress suppressor mutants most relevant to the virulence defect of the Δ*glmR* mutant, we tested all the lysozyme suppressor mutants in a plaque assay. The plaquing assay represents the most complete *ex vivo* assay for virulence of *L. monocytogenes* requiring cellular invasion, cytosolic survival, intracellular replication, and cell to cell spread. In addition to being sensitive to cell wall stress *in vitro*, Δ*glmR* mutants are unable to form wild type sized plaques in fibroblast monolayers (Fig 1A,B). Only second site mutations in *yfhO, gtcA*, and *corA* statistically significantly rescued the Δ*glmR* plaquing defect (Fig. 1B), while second site mutations in *relA*, *pbpA* and *oppA* further inhibited plaquing efficiency of Δ*glmR* mutants. The *yfhO::Tn* and *gtcA::Tn* displayed the most robust suppressor phenotype so we chose to focus on these mutants for follow up studies. Both mutants suppress lysozyme sensitivity, consistent with their identification through the lysozyme suppressor screen (Fig. 1C). In *L. monocytogenes* 1/2a strains, both YfhO and GtcA are required for modification of the WTA repeating ribitol subunits with GlcNAc derived from UDP-GlcNAc (15, 27, 28). We confirmed that the Δ*glmR gtcA::Tn* double mutant is defective for GlcNAc WTA decoration based on loss of wheat germ agglutinin staining (Fig. S1). Finally, disruption of *gtcA* or *yfhO* in a Δ*glmR* mutant partially restores virulence in a murine model of disseminated Listeriosis (Fig. 1D). Taken together, these data suggest that elimination of non-essential decoration of WTA with GlcNAc increases available pools UDP-GlcNAc which can rescue Δ*glmR* mutant cell wall stress sensitivity and virulence ex vivo and in vivo.

**Figure 1.**
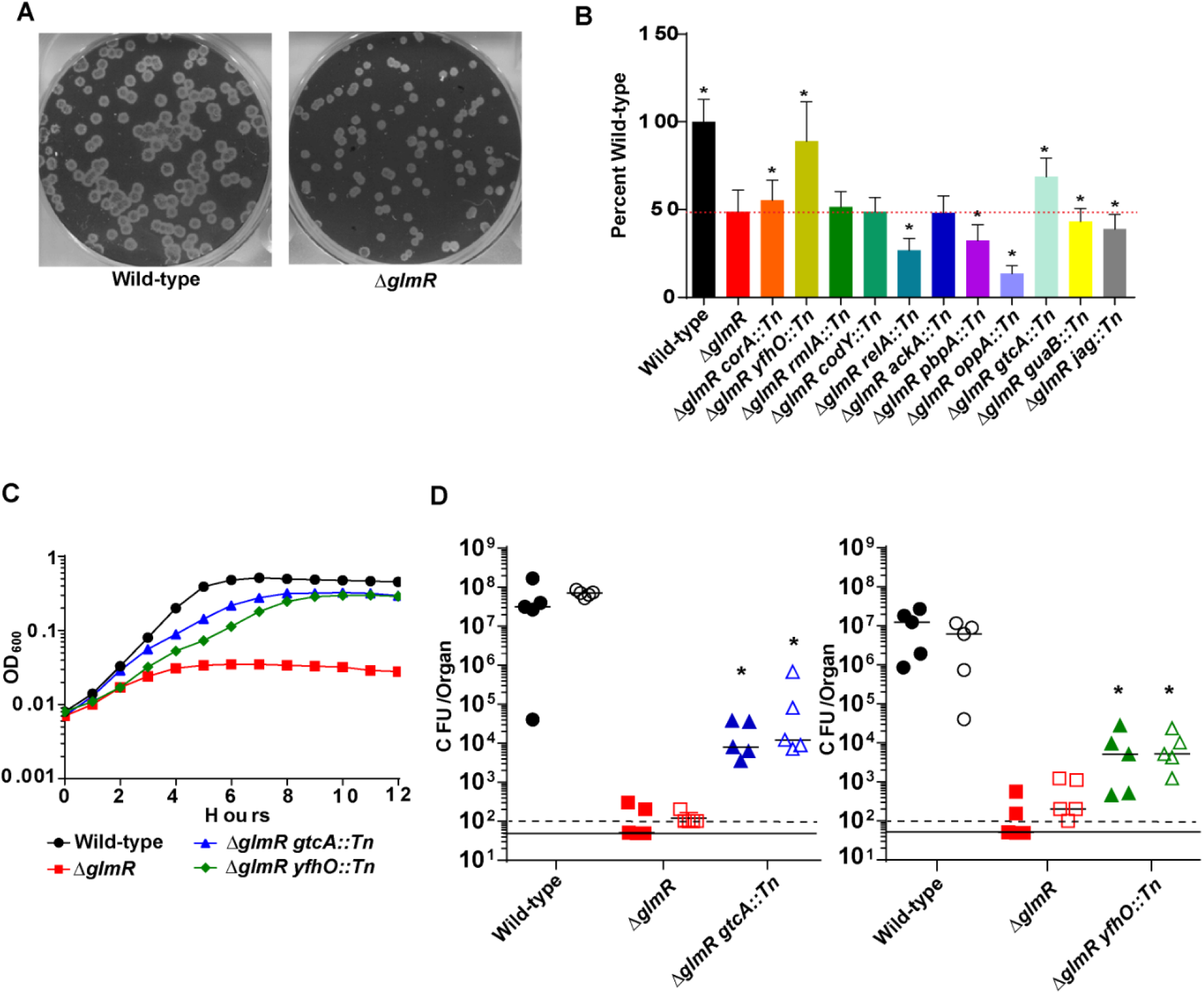
Inhibition of GlcNAc WTA modification suppresses Δ*glmR* mutant phenotypes. **(A)** Representative image of plaques. **(B)** Plaque sizes Δ*glmR* suppressors. Dotted red line indicates Δ*glmR* level. * denotes significant differences from Δ*glmR* by one-way ANOVA (P<0.05). **(C)** Growth in BHI with 1mg/mL lysozyme. Graph is representative of greater than 3 biological replicates. **(D)** CFU from spleens (solid) and livers (open) of C57Bl/6 mice intravenously infected with 1×10^5^ bacteria for 48 hours. The solid line and dotted line represent the limit of detection for spleen and liver respectively. Data are representative of two independent experiments. * denotes significant differences by Mann-Whitney Test (P<0.05)

**Table 1.** ΔglmR suppressor mutants. A Himar 1 transposon mutant library in a Δ*glmR* background was passaged through lysozyme selection. Transposon insertions were identified by sequencing and diagnostic PCR, transduced into a fresh Δ*glmR* background and reconfirmed. Listed are the identified genes, general role they belong to, the number of hits identified in the selection, and the number of unique insertions.

| Table 1 – *ΔglmR* suppressor mutants |  |  |  |  |  |
| --- | --- | --- | --- | --- | --- |
| Role | lmo# | Name | Function | # of hits | #of insertions |
| WTA modification | 1079 | *yfhO* | WTA glycosylation | 9 | 2 |
|  | 1081 | *rmlA* | Glucose-1-phosphate thymidyl transferase | 1 | 1 |
|  | 2549 | *gtcA* | WTA glycosylation | 1 | 1 |
| GTP synthesis and metabolic stress response | 1096 | *guaA* | GMP synthase | 1 | 1 |
|  | 1280 | *codY* | Nutrient response regulator | 1 | 1 |
|  | 1523 | *relA* | ppGpp synthase/reductase | 4 | 2 |
|  | 2753 | *guaB* | Inosine 5'-monophosphate dehydrogenase | 6 | 2 |
| RNA binding | 2853 | *Jag* | Sporulation-related RNA binding protein | 8 | 1 |
| Transport | 1064 | *corA* | Mg transport | 4 | 2 |
|  | 2195 | *oppB* | Oligopeptide ABC transporter | 9 | 1 |
|  | 2196 | *oppA* | Oligopeptide ABC transporter | 15 | 2 |
| Acetate metabolism | 1581 | *ackA* | Acetate kinase | 39 | 1 |
| Peptidoglycan synthesis | 1892 | *pbpA* | High molecular weight penicillin binding protein | 1 | 1 |

### Δ*glmR* mutants have depleted pools of muropeptide precursors

Based on the observation that loss of GlcNAc decoration of the WTA restored lysozyme resistance and partial virulence to Δ*glmR* deficient mutants, we hypothesized that Δ*glmR* mutants may have metabolic defects leading to decreased UDP-GlcNAc synthesis. To test this hypothesis, we utilized untargeted metabolomics to identify differentially abundant metabolites in Δ*glmR* mutants relative to wild type *L. monocytogenes*. After growth in modified *Listeria* synthetic media (LSM) and metabolite extraction, we observed 1073 putative Kyoto Encyclopedia of Genes and Genomes (KEGG) identifiable metabolites including 37 metabolites with >2-fold differences between wild-type and the Δ*glmR* mutant across three biological replicates (Fig. 2A, Table S1). The relatively small number of differential metabolites suggests that GlmR does not have a global regulatory function, at least under laboratory growth conditions. Consistent with our hypothesis, the highest abundance metabolite with >2-fold differential abundance was the essential cell wall precursor metabolite UDP-GlcNAc. Compared to wild type *L. monocytogenes*, UDP-GlcNAc levels are reduced by 73% in the Δ*glmR* mutant (Fig. 2B), consistent with the hypothesis from the suppressor screen that UDP-GlcNAc metabolism is disrupted in the Δ*glmR* mutant. UDP-N-acetyl-muramic acid (UDP-MurNAc), another peptidoglycan precursor downstream of UDP-GlcNAc (Fig. 2C) was similarly decreased in the Δ*glmR* mutant (∼50% of wild type). Upstream of UDP-GlcNAc, N-acetyl-glucosamine-1 phosphate (GlcNAc-1P) levels were also significantly reduced in the Δ*glmR* mutant, however UTP levels were unchanged (Fig. 2B). We were unable to observe the GlmSMU pathway intermediates glucosamine-1 phosphate (GlcN-1P) and glucosamine-6 phosphate (GlcN-6P). Finally, levels of the glycolytic intermediates fructose-6-phosphate (F6P) and fructose-1,6-bisphosphate (FBP) are unchanged in the Δ*glmR* mutant, suggesting that deficits in muropeptide precursors are due specifically to alterations in the GlmSMU pathway and not in a more central metabolic pathway. Levels of UDP-Glucose, a GlmM dependent metabolite, were unchanged between wild-type and the Δ*glmR* mutant indicating that GlmM’s activity is unlikely to be altered in a Δ*glmR* mutant. Consistent with the model that blocking a non-essential UDP-GlcNAc utilizing pathway increases available UDP-GlcNAc for essential PG or WTA synthesis, metabolomic analysis of both the Δ*glmR gtcA::Tn* and the Δ*glmR yfhO::Tn* suppressor mutants demonstrated significant rescue of UDP-GlcNAc levels, though not all the way back to wild-type levels (Fig. 2D). Taken together, these data demonstrate that UDP-GlcNAc metabolism is disrupted in Δ*glmR* mutants and suggests that restoration of UDP-GlcNAc pools restores cell wall stress responses and virulence *in vivo*.

**Figure 2.**
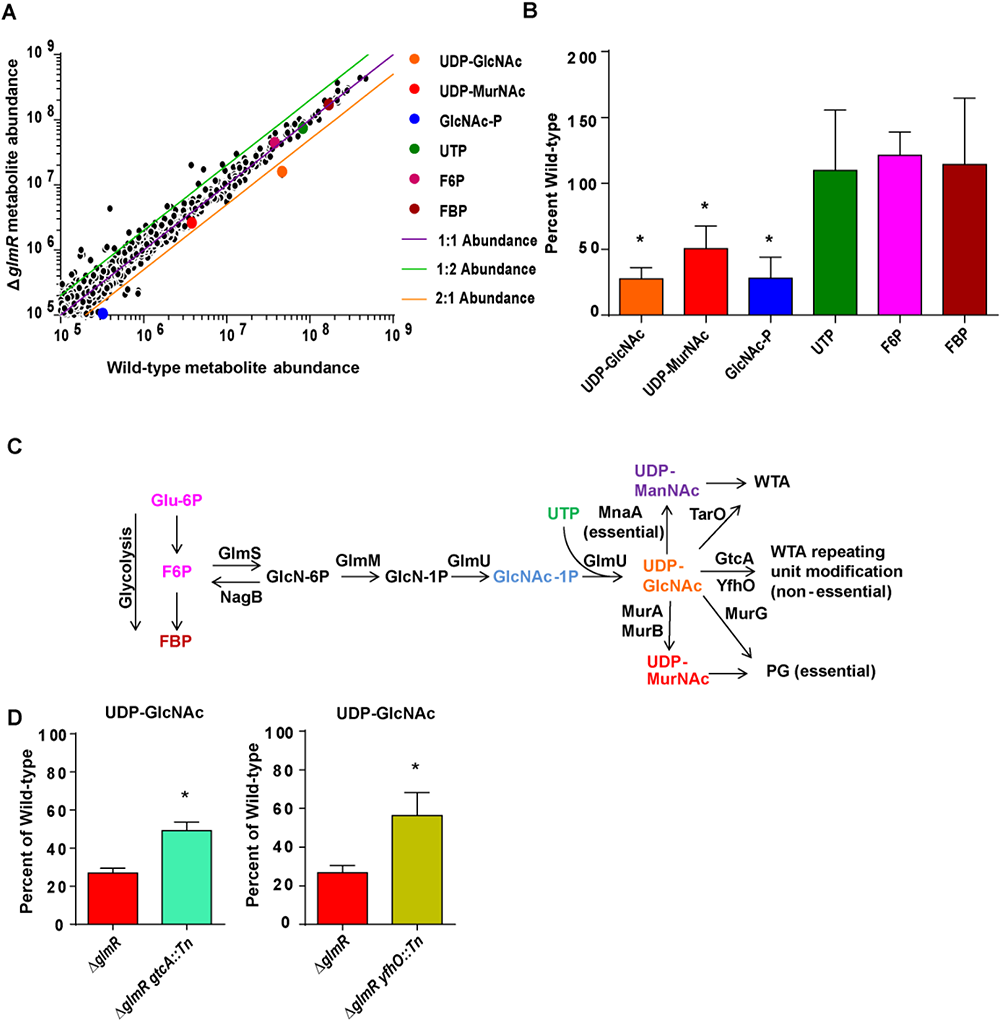
Δ*glmR* mutants are impaired in the production of GlmSMU pathway metabolites. **(A)** Scatter plot of putative KEGG identified ions averaged across 4 biological replicates. **(B)** Quantification of selected metabolites in the Δ*glmR* mutant relative to wild-type across 4 biological replicates. * denotes significant differences from wild-type by student’s t-test (P<0.05). **(C)** UDP-GlcNAc synthesis and utilization pathway **(D)** Quantification of selected metabolites in Δ*glmR* suppressor mutants across 3 biological replicates. * denotes significant differences from Δ*glmR* by student’s t-test (P<0.05).

### GlmR is an accessory uridyltranferase

Two recent studies in *B. subtilis* suggested that GlmR’s function was to enhance the activity of GlmS through direct GlmR-GlmS interactions. Bacterial two-hybrid assays demonstrated a direct interaction between *B. subtilis* GlmR and GlmS (26) and a subsequent study demonstrated that this interaction modulates GlmS activity (29). To determine if GlmR-GlmS interactions were conserved in *L. monocytogenes*, we expressed both *B. subtilis* and *L. monocytogenes* GlmS and GlmR constructs in the bacterial two hybrid system. Each protein was expressed independently as both N-and C-terminal fusions to both T18 and T25. Four replicates of the blue-white assay were performed due to variability in the system from a known thresholding effect (30) and quantitative β-galactosidase assays were performed in triplicate. As predicted based on their crystal structures, GlmS (31) and GlmR (PDB 2Q7X and 1HZB) from both *B. subtilis* and *L. monocytogenes* homodimerized, demonstrating that the constructs were expressed and functional (Fig S2A,B). Positive, but inconsistent interactions between *B. subtilis* GlmR and GlmS were observed as previously reported for one set of *B. subtilis* fusion proteins (Fig S3A,B) (26), however no combination of *L. monocytogenes* GlmR and GlmS produced an interaction except those for which there was also activity observed in the empty vector controls (Fig S4A,B). Taken together these data suggest that GlmR regulation of GlmS through protein-protein interactions may not be evolutionarily conserved among GlmR homologues and that GlmR must function to regulate UDP-GlcNAc levels by a novel mechanism in *L. monocytogenes*.

A distant homologue of GlmR is CofD, a 2-phospho-l-lactate transferase involved in the synthesis of Coenzyme F420 in actinobacteria (32). This homology to a catalytic protein suggests that perhaps GlmR has direct enzymatic activity. We hypothesized that, analogous to the accessory UDP-N-acetylglucosamine enolpyruvyl transferase function of MurZ (33), GlmR could be an accessory enzyme functioning to increase pools of UDP-GlcNAc in the context of cell wall stress. To test this hypothesis, we cloned and purified GlmR from *L. monocytogenes* and assessed its potential enzymatic activity in the last two steps of the GlmSMU normally catalyzed by GlmU to produce UDP-GlcNAc. Using mass spectrometry to assess the results of each reaction, we found that GlmR catalyzed the synthesis of UDP-GlcNAc from GlcNAc-1P and UTP (Fig 3A,B), similar to both commercially purchased *Escherichia coli* GlmU as well as *L. monocytogenes* GlmU that we expressed and purified (Fig. 3A,B). Importantly, no UDP-GlcNAc was detectable with substrates UTP and GlcNAc-1P alone indicating that catalysis required either the GlmU or GlmR protein (Fig. 3B). Additionally, purified GlmR demonstrated no acetyltransferase activity, demonstrating that the activity observed was not an artifact of accidental co-purification of GlmU (Fig. S5A). Finally, the absence of UDP-GlcNAc in a GlmR reaction mixture lacking GlcNAc-1P and UTP as substrates or after the protein was heated excludes the possibility of UDP-GlcNAc being a co-purified artifact with GlmR (Fig. 3B). Taken together, these data suggest that GlmR can act as a uridyltransferase enzyme to facilitate increased production of UDP-GlcNAc in response to cell wall stress.

**Figure 3.**
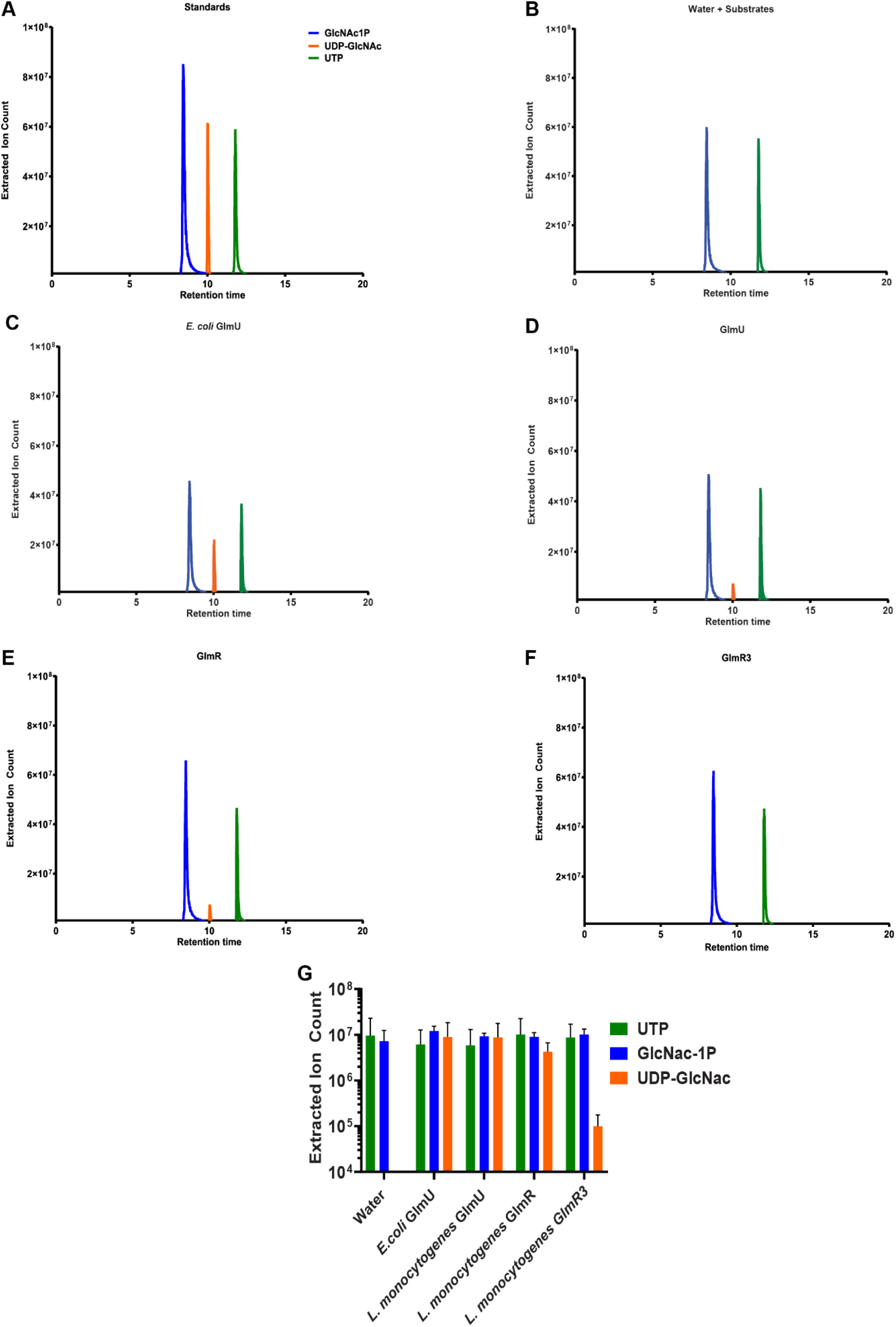
GlmR catalyzes the production of UDP-GlcNAc. **(A)** HPLC-MS analysis of reactions with 100μM substrates alone or in combination with 1μM purified GlmU or GlmR as indicated. Extracted Ion Counts for the relevant metabolites are indicated based on purified standards (GlcNAc-1P blue, UTP green, UDP-GlcNAc orange). **(B)** Quantification of selected metabolites (GlcNAc-1P blue, UTP green, UDP-GlcNAc orange) from reactions with 100µM substrates alone or in combination with water, 1µM *E.coli* GlmU, GlmU, GlmR, and GlmR3. Assays were performed in triplicate.

### GlmR uridyltransferase activity is conserved

GlmR is the second gene of a highly conserved operon of three genes found in firmicutes and actinobacteria. In *S. aureus* the GlmR homologue YvcK is predicted to be essential (21) while in *M. tuberculosis* the GlmR homologue CuvA is essential for virulence (19, 20). The *S. aureus* and *M. tuberculosis* homologues from these species exhibit high identity to *L. monocytogenes* GlmR, with 46% identity, 69% similarity and 34% identity, 57% similarity, respectively, and are best conserved near the putative N-terminal active site (Fig. 4A). To determine whether GlmR enzymatic function is broadly conserved, we first purified GlmR from *S. aureus*, *M. tuberculosis* and *B. subtilis* and assessed enzymatic activity. Each protein exhibited uridyltransferase activity similar to *L. monocytogenes* GlmR (Fig. 4B). To test for functional conservation of GlmR function *in vivo* we complemented the *L. monocytogenes* Δ*glmR* mutant with codon optimized *glmR* homologues from *S. aureus* and *M. tuberculosis.* Despite the high sequence conservation and the conserved enzymatic activity *in vitro*, only the *S. aureus* homologue was able to complement lysozyme sensitivity (Fig. 4C). The *B. subtilis* GlmR homologue was also able to rescue lysozyme resistance of *L. monocytogenes* Δ*glmR* mutant (Fig S6). Consistent with their ability to rescue lysozyme resistance, we found that complementation of the *L. monocytogenes ΔglmR* mutant with the *S. aureus* homologue but not the *M. tuberculosis* homologue restored UDP-GlcNAc levels (Fig. 4D). Taken together, these data suggest that the uridyltransferase enzymatic function of GlmR is highly conserved, however the observation that *M. tuberculosis* CuvA cannot transcomplement an *L. monocytogenes* Δ*glmR* mutant suggests additional mechanisms of GlmR regulation including potentially protein localization and/or post-translational modification (16, 19, 34).

**Figure 4.**
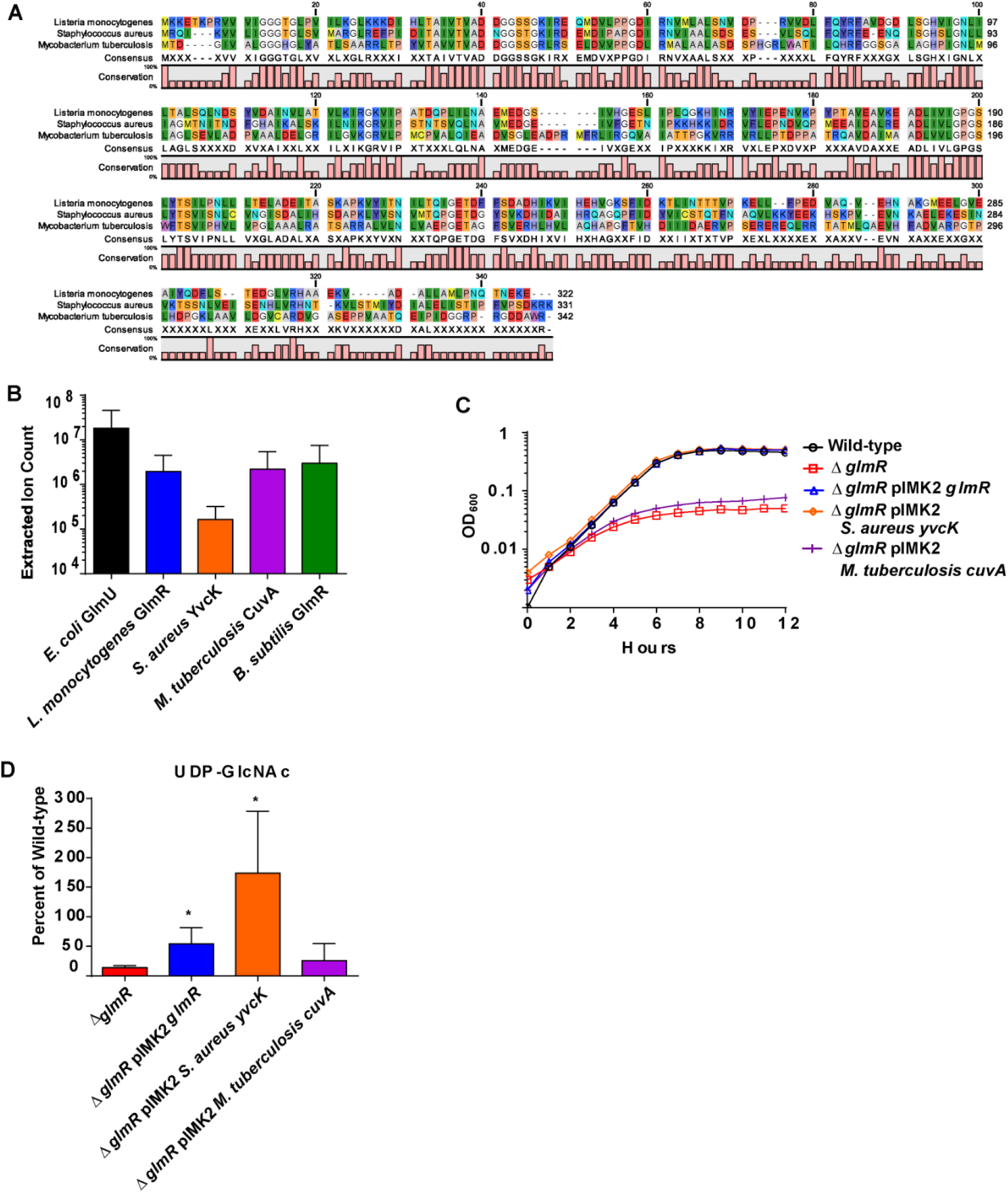
GlmR uridyltransferase function is conserved in *S. aureus* and *M. tuberculosis*. **(A)** GlmR homologues aligned using CLC Sequence Viewer 8.0. **(B)** Analysis of uridyltransferase activity of *E. coli* GlmU and purified GlmR homologues by HPLC-MS. No significant differences by ANOVA. **(C)** Transcomplementation of growth in BHI with 1mg/mL lysozyme over 12 hours at 37°C. Graph is representative of greater than 3 biological replicates. **(D)** Transcomplementation of UDP-GlcNAc levels relative to wild-type. * denotes significant differences from *glmR* by student’s t-test (P<0.05).

### GlmR uridyltransferase activity is required for cell wall stress responses and virulence *in vivo*

Our data suggested that GlmR can act as an accessory uridyltransferase. Based on modeling against the crystal structure of the homologue PDB:2O2Z (Fig. 5A), we predicted D40, D41 and N198 to be active site residues that when mutated to alanines would abolish catalytic activity. To test the hypothesis that uridyltransferase activity is necessary for virulence we created a D40A D41A N198A mutant GlmR (GlmR3), purified the mutant protein and assessed uridyltransferase activity. Activity of the GlmR3 mutant was ∼100-fold reduced in an in vitro biochemical assay compared to wild type GlmR (Fig. 3A,B). Complementation of a *glmR* mutant with *glmR3* was unable to rescue lysozyme sensitivity (Fig 5B) despite equal or even increased levels of expression compared to the wild type GlmR complement (Fig. S7) Finally to test the hypothesis that uridyltransferase activity is important for virulence we performed infected mice and quantified bacterial burdens in an *in vivo* model of disseminated Listeriosis. In contrast to complementation with wild type GlmR, the GlmR3 mutant was unable to rescue the virulence defect of the Δ*glmR* mutant (Fig. 5C). Taken together, these data suggest that the uridyltransferase activity of GlmR is essential for mediating cell wall stress responses during infection to facilitate virulence of *L. monocytogenes*.

**Figure 5.**
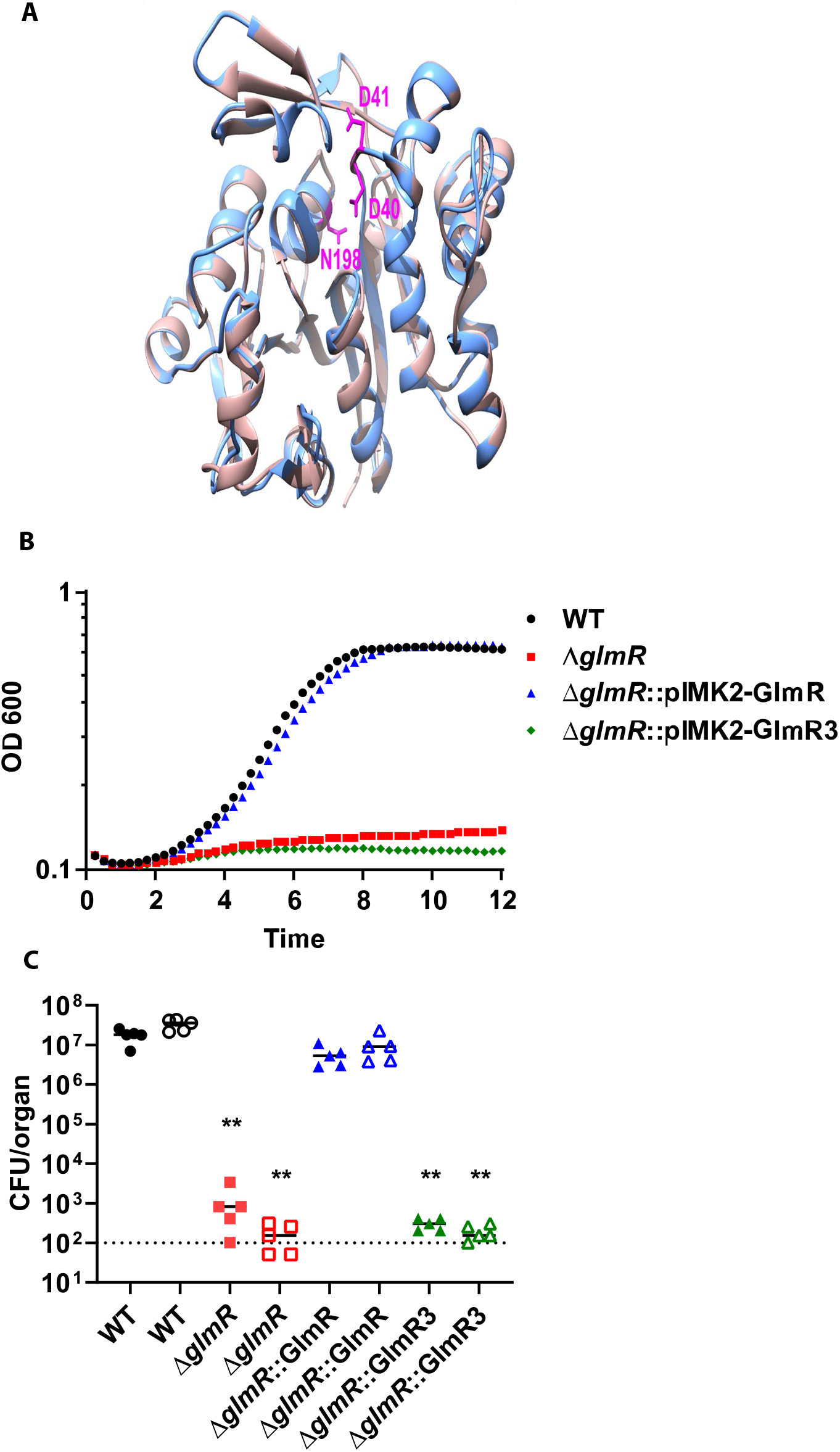
GlmR uridyltransferase activity necessary for virulence. **(A)** The amino acid sequence of *L. monocytogenes* GlmR(light blue) is highly similar to that of *Bacillus halodurans* homolog (light pink) (∼47% sequence identity, for which the crystal structure (PDB 2O2Z) has been solved. Based on this similarity, we used Phyre2 to generate an *Lm* GlmR structural model, using the 2O2Z structure as a template, and found the two structures to be superimposable. Mutations made in the predicted catalytic site are highlighted (hot pink) D41, D40, N198 (from top, clock-wise). **(B)** Growth of WT, Δ*glmR*, Δ*glmR-pIMK2-GlmR*, and Δ*glmR: pIMK2-GlmR3* in BHI with 1 mg/mL of lysozyme over 12 hours at 37°C. Graph is representative of greater than 3 biological replicates. **(C)** C57/Bl6 mice were infected intravenously with 1×10^5^ wild-type (black circles), Δ*glmR*mutants (red squares), Δ*glmR::GlmR* mutants (blue triangles), Δ*glmR::GlmR*3 (green triangles) *in vivo*. Spleens (solid) and liver (open) were harvested 48 hours post infection homogenized and plated for CFU. The median (solid bar) and limit of detection (dotted line) for each experiment indicated. Data are representative of two independent experiments with 5 mice each. * indicates statistical significance by Mann-Whitney test (P<.05).

## Discussion

GlmR is a highly conserved protein that is essential for virulence in *L. monocytogenes* and *M. tuberculosis*, but whose function remains largely unknown (16, 19, 20). In this study we discovered that GlmR has conserved uridyltransferase activity that facilitates cell wall stress responses during infection. Our findings are also consistent with a recent study utilizing *t*-Cin hypersensitive *L. monocytogenes glmR*:Himar1 mutants which identified suppressor mutations in genes involved in the biosynthesis of UDP-GlcNAc (53). When the *glmR*: Himar1 mutant was engineered to overexpress *glmU* growth in t-Cin was fully restored, whereas overexpression of *glmS*, or *glmM* only partially restored resistance to *t*-Cin further supporting the idea that GlmR is involved in the biosynthesis of UDP-GlcNAc and that the terminal step of the GlmSMU pathway is rate limiting (53). Deciphering the activities of proteins of unknown function, such as GlmR, is a major challenge not only in microbial pathogenesis but in biology at large. Indeed, 25% of predicted biochemical reactions do not have an assigned enzyme, suggesting that many proteins of unknown function have enzymatic activity (35, 36). Recent metabolomics approaches such as activity-based metabolomics have shown great promise in identifying these functions (36, 37). Combining parallel screening approaches such as genetics, transcriptomics, proteomics, and metabolomics generates targeted hypotheses about the roles of proteins of unknown function in physiological processes. In this study an untargeted metabolomics approach combined with a classical bacterial genetics suppressor screen allowed us to discover the uridyltransferase activity possessed by GlmR.

Although GlmR has potentially separable functions in both central metabolism and cell wall homeostasis (22), our identification of suppressor mutations that rescue virulence through restoration of UDP-GlcNAc levels suggests that GlmR’s role in cell wall homeostasis is critical during infection. GlmR’s function in promoting cytosolic survival further suggests that bacteria experience cell wall stress in the cytosol, however the cytosolic CAD responsible for imparting cell wall stress is unknown. Guanylate Binding Proteins (GBPs) and Lysozyme are not responsible for the cytosolic cell wall stress as GlmR is required for cytosolic survival even in Gbp^Chr3-/-^ and LysM^-/-^ macrophages (16, 38). Future identification of the cytosolic CADs targeting the bacterial cell wall will illuminate novel host defense pathways, not only against *L. monocytogenes*, but also other bacteria that invade the cytosol, including both canonical and non-canonical cytosolic pathogens such as *M. tuberculosis* and *S. aureus*. Furthermore, other bacterial pathogens which require GlmR for survival and virulence, such as *S. aureus* (21) and *M. tuberculosis* (19, 20), likely require GlmR to deal with cell wall stress in their conventional replication niches.

We found that GlmR uridyltransferase activity is conserved in *S. aureus* and *M. tuberculosis*, representatives of the Firmicutes and Actinobacteria phyla. This conservation combined with its essential role in virulence of a number of important pathogens suggest that it may be an attractive drug candidate. Indeed, both the acetyl-and uridyltransferase activities of *M. tuberculosis* GlmU have been targeted by small molecules as a novel antibiotic strategy (39). Whether uridyltransferase inhibitors of GlmU could also bind and inhibit GlmR will need to be assessed. Among GlmR homologues, the N-terminal putative nucleotide binding region is most highly conserved. This raises important questions not only about the design of GlmR small molecule inhibitors, but also about substrate specificity of GlmR homologues and whether different GlmR proteins may have flexibility to catalyze different reactions with regard to the sugar component. Indeed, this may explain why GlmR appears to have separable roles in both cell wall homeostasis and gluconeogenic metabolism. Crystal structures of GlmR homologues in complex with their substrates will be critical both for antibiotic development and an understanding of the potential promiscuity of these enzymes.

GlmR uridyltransferase activity is conserved, but the ability to transcomplement *L.monocytogenes* Δ*glmR* mutants is not, suggesting that regulation of GlmR activity is essential. In *L. monocytogenes*, GlmR is upregulated at the protein level by cell wall stress (16), but the underlying mechanism of this upregulation remains unknown. Additionally, GlmR is phosphorylated by PASTA kinases in *L. monocytogenes*, *B. subtilis*, and *M. tuberculosis*, however the phosphorylation sites differ and what effect phosphorylation may have on the enzymatic activity is similarly unknown (16, 19, 40). Subcellular localization of GlmR may also contribute to its regulation as GlmR localization patterns in *B. subtilis* and *M. tuberculosis* are dissimilar (19, 22, 34, 41). Finally, recent studies suggested that GlmR may also act allosterically to regulate the function of GlmS in *B. subtilis* (26, 29). Although we were unable to observe this interaction in *L. monocytogenes*, GlmR functioning as an allosteric regulator of GlmS and as a functional uridyltransferase are not mutually exclusive and indeed could act synergistically. Identification of mutations which abolish GlmS-GlmR interaction but not enzymatic activity and vice versa are necessary to separate and test these ideas. This study identified that GlmR, a protein required for *L. monocytogenes* and *M. tuberculosis* virulence, is an accessory uridyltranferase necessary for UDP-GlcNAc synthesis in the context of cell wall stress. Similar to MurA and MurZ in *S. aureus* (33), this highlights that virulence determinants can be redundant with essential housekeeping enzymes. Often these accessory enzymes are upregulated in the context of stress, such as during infection or antibiotic treatment as is the case with GlmR and MurZ, respectively (33). Indeed GlmR’s enzymatic activity may have gone previously undiscovered despite its importance due to the protein’s low expression during normal laboratory culture with rich media. Additionally, with a potential exception in *S. aureus* (21), GlmR is likely not essential under laboratory conditions due to sufficient uridyltransferase activity of GlmU. Conversely, even in a situation where GlmR complemented GlmU uridyltransferase activity, GlmU would still be essential due to its acetyl-transferase function. Future analysis of virulence determinants of unknown function through parallel screening approaches may reveal this redundancy to be even more pervasive.

## Methods

### Listeria monocytogenes strains and culture

All *L. monocytogenes* strains used for experiments in this study were 10403S background. The Δ*glmR* mutant was described previously (14). *L. monocytogenes* was grown overnight in BHI at 30°C for all experiments except as described for metabolomic analysis.

### Construction of *L. monocytogenes* strains

Homologue complementation genes used in Figure 4 were created with gBlocks (IDT) that were codon-optimized for *L. monocytogenes* and inserted into pIMK2 (42) under control of the constitutive pHelp promoter. The complementation construct used in Fig S4 was not codon-optimized and inserted into pPL2e under control of a theophylline inducible riboswitch as previously described (43). Constructs were cloned in XL1-Blue *E. coli* with 30μg/mL Kanamycin for pIMK2 and 2μg/mL Erythromycin for pPL2e as appropriate and shuttled into *L. monocytogenes* through conjugation with SM10 or S17 *E. coli*.

### Suppressor Selection

A Himar 1 Tn mutant library was created in a Δ*glmR* mutant background as described previously (44). Aliquots of library were thawed, diluted 1:1000-10000 in PBS and inoculated 1:50 into 1mL of LB with 1mg/mL lysozyme and 0.1uM staurosporine in pentaplicate. 50μL of cultures were plated at 0 hours on LB and 6 hours on LB 1mg/mL lysozyme. This selection was carried out four times and 313 out of 476 resulting colonies were secondarily screened in BHI with lysozyme 1mg/mL staurosporine 0.1μM. Transposon mutations in the remaining suppressors were identified by 2-step PCR using transposon specific and degenerate primers followed by sanger sequencing using and were confirmed by PCR with diagnostic primers (45). To determine whether identified transposon mutations were causative, all unique transposons were transduced into a fresh Δ*glmR* background and reconfirmed with diagnostic PCR, sequencing, and rescue of the Δ*glmR* mutant lysozyme sensitivity with overnight growth in 1mg/mL lysozyme in BHI.

### Phage Transduction

Phage transductions were performed as previously described (46). Briefly, U153 phage stocks were propagated with MACK *L. monocytogenes* grown overnight in LB at 30°C. MACK cultures were pelleted and resuspended in LB with 10mM MgCl2 and 10mM CaSO4 and mixed with 0.7% LB agar 10mM MgCl2 10mM CaSO4 at 42°C and immediately poured on LB plates and incubated overnight at 30°C. Plaque lysate was soaked out with 10mM Tris pH 7.5 10mM MgCl2 10mM CaSO4 buffer, and sterilized by 0.2μm filtration or addition of 1:3 volume chloroform. Donor plaque lysates were prepared using the same conditions and used to infect recipient Δ*glmR* cultures for 1 hour at room temperature before being plated on erythromycin selection at 37°C.

### Lysozyme Sensitivity

Overnight 30°C static BHI cultures were backdiluted 1:50 into 96-well plates containing BHI or BHI with lysozyme at 1mg/mL. Plates were grown at 37°C with continuous shaking for 12 hours in an Eon or Synergy HT Microplate Spectrophotometer (BioTek Instruments, Inc., Winooski, VT) and OD_600_ was read every 15 minutes.

### Plaque Assay

The plaque assay was performed as described (47) except that the MOI was adjusted for optimal plaque number and an additional plug was added to wells at 3 days to facilitate an additional 3 days of plaque growth. At 6 days wells were stained with 0.3% crystal violet and washed with water. After staining the dishes were scanned and plaque areas were quantified with ImageJ. All strains were assayed in biological triplicate and the plaque areas of each strain were normalized to wild-type plaque size within each replicate.

### Metabolite Extraction

Overnight 30°C static BHI cultures were washed with PBS and backdiluted 1:50 into 50mL of Listeria synthetic media (LSM) baffled flasks 37°C shaking and grown to an OD600 of ∼0.4. LSM is a derivative of Improved Minimal Media developed by Phan-thanh and Gorman (48) with several component changes (49). For metabolomic experiments we reduced the level of MOPS to 1/5^th^ the normal amount to reduce background MS signal. 5mL of culture was deposited by vacuum filtration onto a 0.2 µm nylon membrane (47 mm diameter) in duplicate. The membrane was then placed (cells down) into 1.5 ml cold (−20°C or on dry ice) extraction solvent (20:20:10 v/v/v acetonitrile, methanol, water) in a 60mm petri dish and swirled. After a few moments the filter was inverted (cells up) and solvent was passed over the surface of the membrane several times to maximize extraction. Finally, the cell extract was stored at −80°C. Extracts were pelleted at 21000 rcf at 4°C for 10 minutes. ∼200μL of extract normalized to OD was dried with N2 gas. Extracts were resuspended in 70μL of HPLC grade water and pelleted at 21000 rcf at 4°C for 10 minutes to remove particulates. All cultures were extracted in biological triplicate or quadruplicate and in technical duplicate.

### Metabolite quantification and analysis

Metabolite quantification and analysis was performed with the same instrument and chromatography set up as previously described (50). Briefly, samples were run through an ACQUITY UPLC® BEH C18 column in a 18 minute gradient with Solvent A consisting of 97% water, 3% methanol, 10 mM tributylamine (TBA), 9.8 mM acetic acid, pH 8.2 and Solvent B being 100% methanol. Gradient was 5% Solvent B for 2.5 minutes, gradually increased to 95% Solvent B at 18 minutes, held at 95% Solvent B until 20.5 minutes, returned to 5% Solvent B over 0.5 minutes, and held at 5% Solvent B for the remaining 4 minutes. Ions were generated by heated electrospray ionization (HESI; negative mode) and quantified by a hybridquadrupole-high-resolution mass spectrometer (Q Exactive orbitrap, Thermo Scientific). MS scans consisted of full MS scanning for 70-1000 *m/z* from time 0–18 min except MOPS m/z of 208-210 was excluded from 1.5-3 minutes. Metabolite peaks were identified using Metabolomics Analysis and Visualization Engine (MAVEN) (51, 52).

### Protein Purification

#### GST tagged expression and purification scheme

GlmR, GlmR3 and GlmU were cloned into pGex6P into XL1-Blues and expressed in Rosettas with pLysS. IPTG was added to 500 uM to induce expression and 3 hours post induction cells were pelleted, resuspended in PBS, and frozen at −80C. Cell suspensions were thawed and lysed by sonication in the presence of protease inhibitors. Cell debris was pelleted and cell lysate was filtered with a 0.2 μM filter and loaded onto a prepacked glutathione resin column at 4°C. The column was washed two times with 10 column volumes of cleavage buffer (25 mM Tris pH8 100 mM NaCl 1mM DTT) before elution. The column was loaded with 80 units of PreScission Protease in 960 uL of cleavage buffer and incubated overnight at 4°C. Elution was collected the next day by adding 3 mL of cleavage buffer to the column and concentrated between 15 uM and 23 uM. Protein was stored at 4°C and purity was assessed by SDS-Page and protein was quantified by BCA assay.

#### Hist-Tagged expression and purification scheme

GlmR homologues were cloned into pET20b in XL1-blues and expressed in Rosettas with pLysS except for CuvA which was expressed from BL-21s from pET23. IPTG was added to 500uM to induce expression and 3 hours post induction cells were pelleted, resuspended in PBS, and frozen at −80°C. Cell suspensions were thawed and lysed by sonication in the presence of protease inhibitors. Cell debris was pelleted and cell lysate was filtered with a 0.2μm filter and loaded onto a HisTrap Ni column (GE Healthcare) at 4°C. The column was washed with PBS and PBS 25mM Imidizole before elution with 250mM Imidizole. Elutions were dialyzed overnight at 4°C into 10mM Tris pH 7.4 100mM NaCl which was prepared at 25°C and concentrated to between 6 and 22μM. Protein was stored at 4°C and purity was assessed by SDS-PAGE and protein was quantified by BCA assay.

### Enzymatic Activity

Reactions were carried out in 10mM Tris pH 7.4, 100mM NaCl, 1mM MgCl2 buffer. Substrates were added at 100uM and purified *E. coli* GlmU (Galen Laboratory Supplies, GL01012), *L. monocytogenes* GlmU, *L. monocytogenes* GlmR or GlmR homologues were added at 1μM and incubated at 37°C for 10 minutes. Protein was removed with a 3kDa MWCO filter, resulting reaction mixtures were diluted 1 to 10 in solvent A and analyzed by tandem HPLC-MS and Maven software.

### Bacterial Two-Hybrid

GlmR and GlmS from both *L. monocytogenes* and *B. subtilis* were cloned in-frame into vectors pU18, pU18C, pKT25, and pKNT25 from the BACTH System Kit (Euromedex) using XbaI and KpnI. Constructs were made originally in TAM1 or XL1-Blue *E. coli* and then moved to BTH101 *E. coli* for testing. Both blue/white screening on X-gal plates and β-Galactosidase assays were carried out as previously described (30).

### Mouse infection

Infections were performed as previously described (16). Briefly, 6 to 8-week-old female and male C57BL/6 mice were infected IV with 1×10^5^ CFU. 48 hours post-infection, livers and spleens were harvested, homogenized in PBS with 0.1% NP-40, and plated for CFU. Two independent replicates of each experiment with 5 mice per group were performed.

### Ethics statement

Mice were cared for according to the recommendations of the NIH, published in the Guide for the Care and Use of Laboratory Animals. All techniques used were reviewed and approved by the University of Wisconsin-Madison Institutional Animal Care and Use Committee (IACUC) under the protocol M005916.

### Statistical Analysis

Prism 6 (GraphPad Software) was used for statistical analysis of data. Means from two groups were compared with unpaired two-tailed Student’s T-test. Means from more than two groups were analyzed by one-way ANOVA with a post-hoc LSD Test. Mann-Whitney Test was used to analyze non-normal data from animal experiments. * indicates a statistically significant difference (P is less than 0.05).

## Acknowledgements

This work was supported by R01 AI137070 (JD.S.) and R01 AI097157 (J.P.D.).

## Supplementary Information

### Supplementary Methods

#### Wheat Germ Agglutinin Staining

1mL of overnight cultures in BHI at 37°C were pelleted, fixed in 4% paraformaldehyde in PBS, washed in PBS with 0.1% Tween (PBS-T), resuspended in 100μL PBS-T, and incubated with 50μL of 0.1% Wheat Germ Agglutinin (WGA) for 5 minutes. Pellets were washed in PBS-T and stored at 4°C in the dark. Confocal microscopy was performed as previously described (43).

---

**Figure S1.**
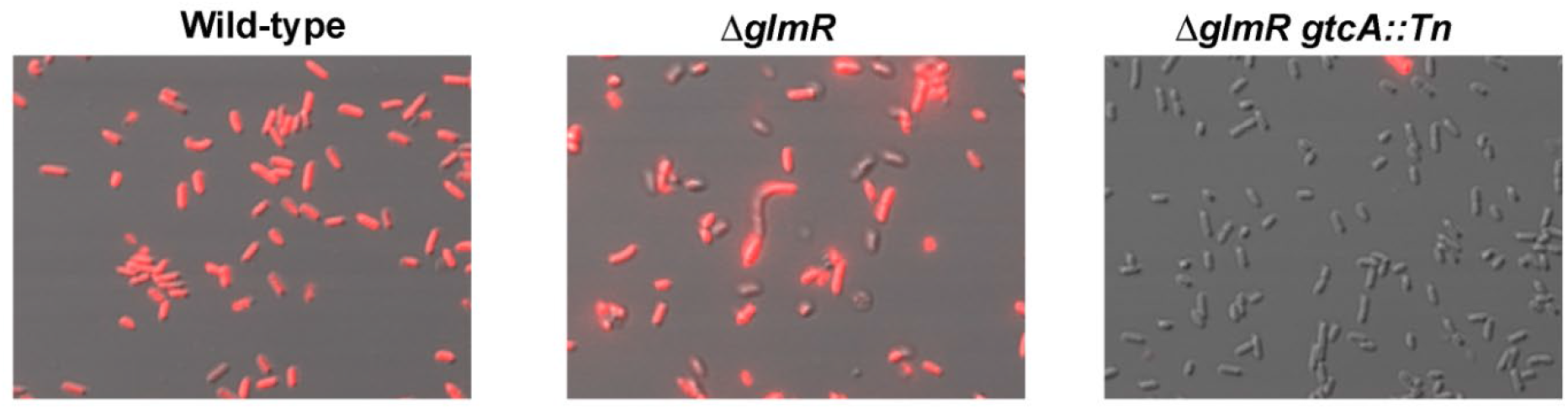
GtcA is functionally inactivated by a Tn insertion. Wild-type, *glmR*, and *glmR gtcA::Tn* strains were imaged and assessed for their ability to bind WGA (red).

**Figure S2.**
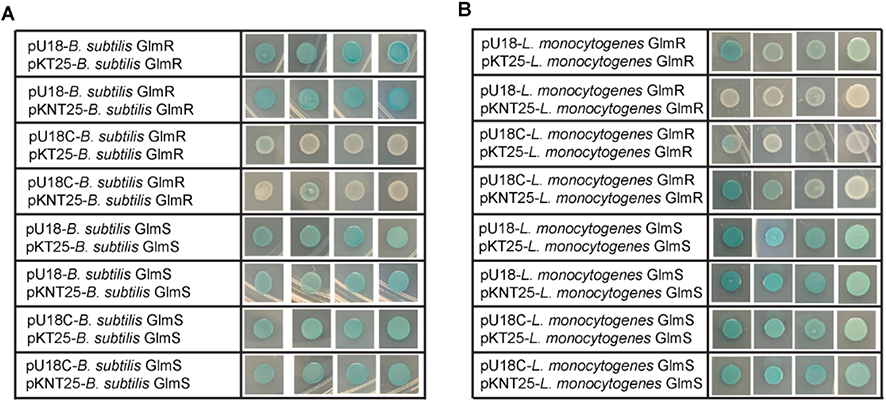
GlmS and GlmR form homodimers. **(A,B)** Bacterial 2-hybrid strains were plated on X-Gal and incubated for 24 hours at 30°C in biological quadruplicate.

**Figure S3.**
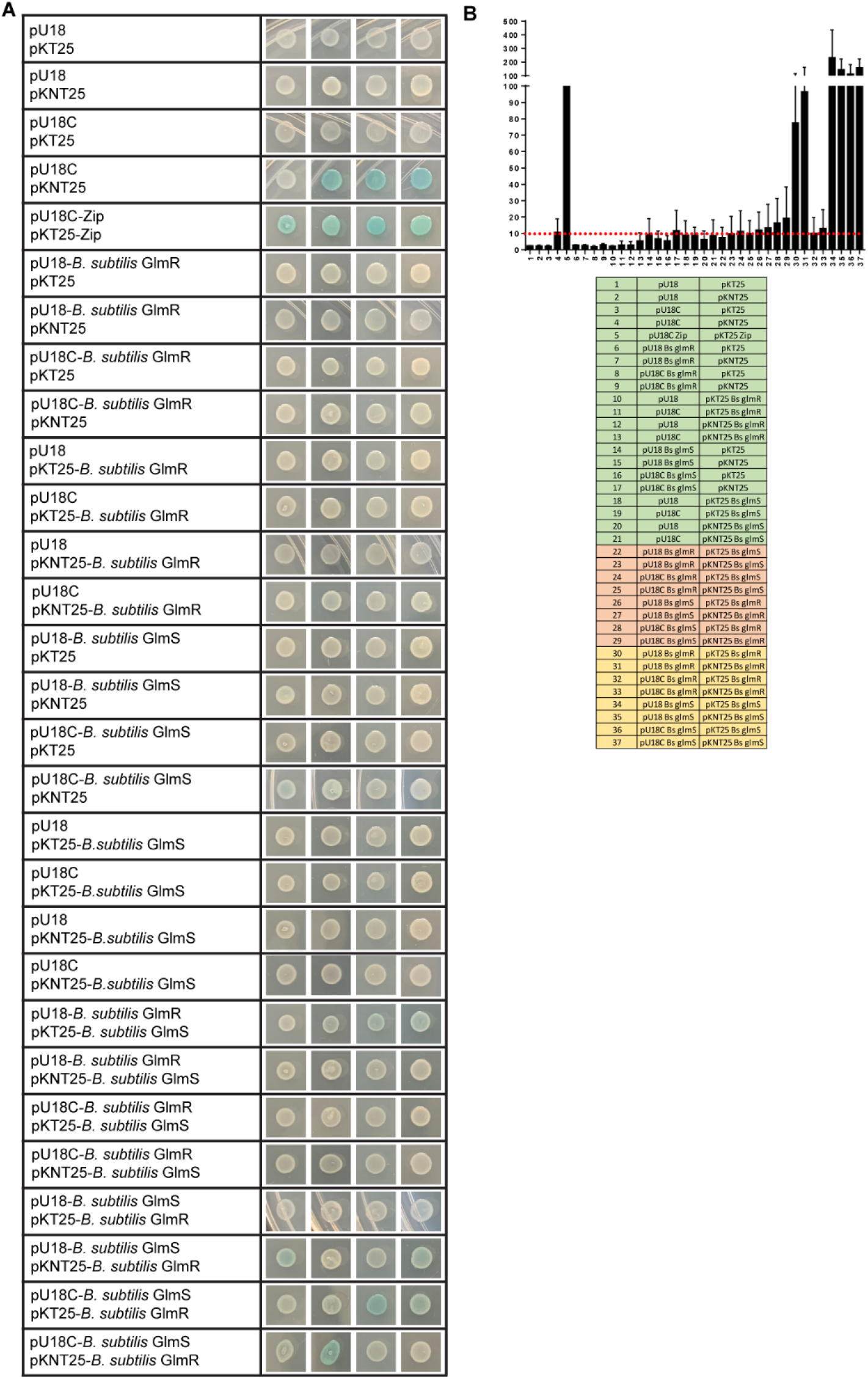
*B. subtilis* GlmR interaction with GlmS. **(A,B)** Bacterial 2-hybrid strains were plated on X-Gal and incubated for 24 hours at 30°C in biological quadruplicate. Bacterial 2-hybrid cultures were lysed and assayed for β-galactosidase activity in biological triplicate. Activity is normalized to the Zip positive control. The dotted red line indicates 10% of the Zip value. Strains are identified by a number and listed below. Control strains are green, GlmS-GlmR interaction test strains are red, and GlmS or GlmR homodimer strains are gold.

**Figure S4.**
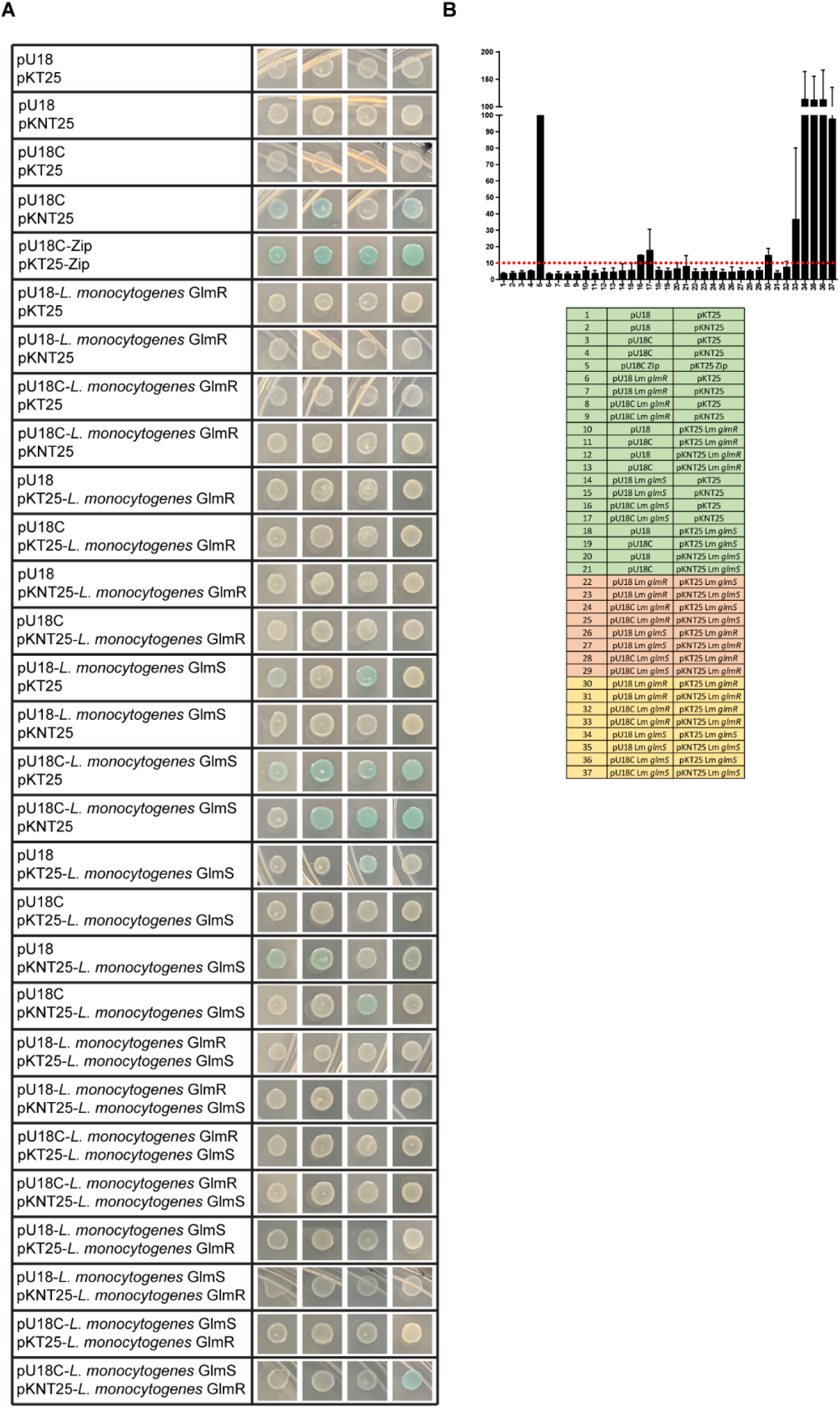
*L. monocytogenes* GlmR does not interact with GlmS. **(A) (B)** Bacterial 2-hybrid strains were plated on X-Gal and incubated for 24 hours at 30°C in biological quadruplicate. Bacterial 2-hybrid cultures were lysed and assayed for β-galactosidase activity in biological triplicate. Activity is normalized to the Zip positive control. The dotted red line indicates 10% of the Zip value. Strains are identified by a number and listed below. Control strains are green, GlmS-GlmR interaction test strains are red, and GlmS or GlmR homodimer strains are gold.

**Figure S5.**
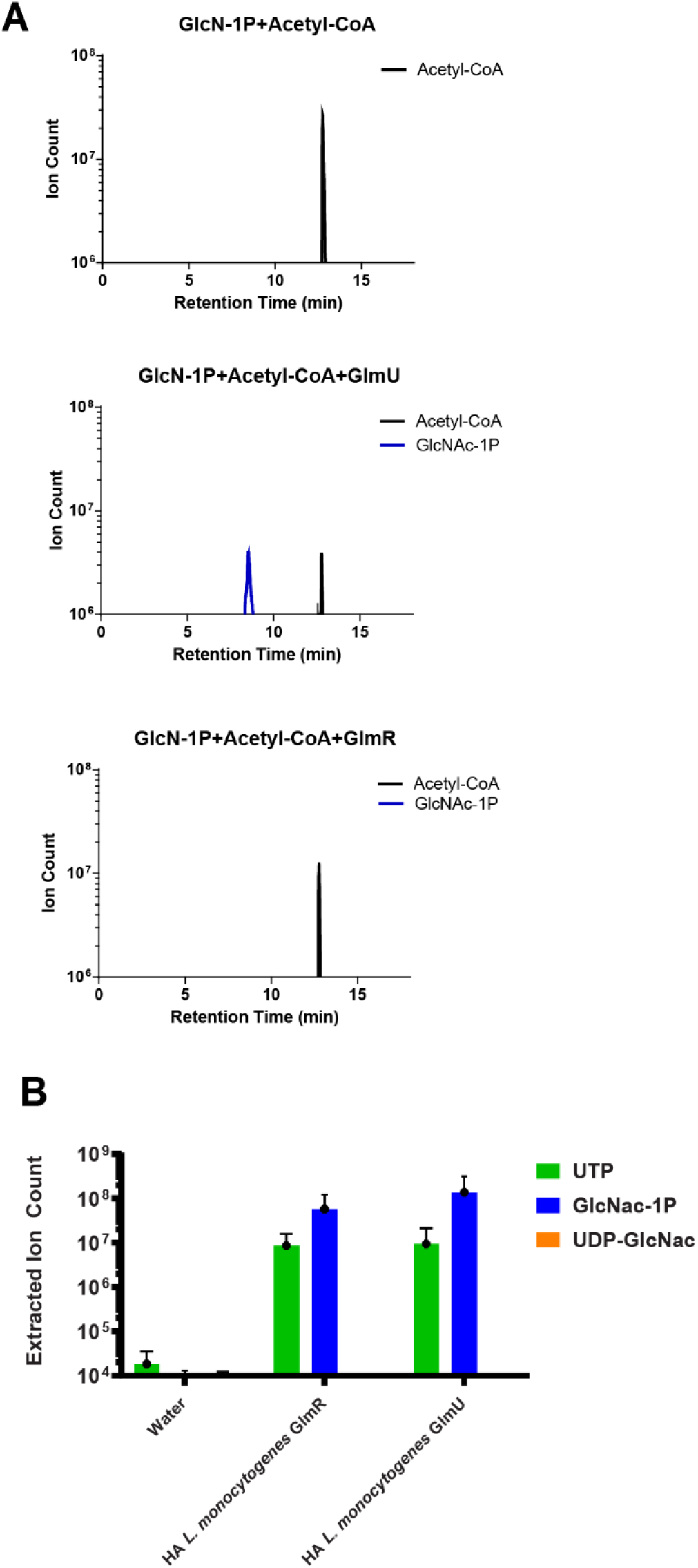
*L. monocytogenes* GlmR lacks acetyltransferase activity. HPLC-MS analysis of reactions with 100μM substrates alone or in combination with 1μM GlmU or GlmR. Peaks for the relevant metabolites are indicated (Acetyl-CoA black, GlcNAc-1P blue, UDP-GlcNAc orange).(B) Quantification of selected metabolites (GlcNAc-1P blue, UTP green, UDP-GlcNAc orange) from reactions with 100µM substrates alone or in combination with water, 1µM Heat Inactivated (HA) GlmR or Heat Inactivated (HA) GlmU. Assays were performed in triplicate.

**Figure S6.**
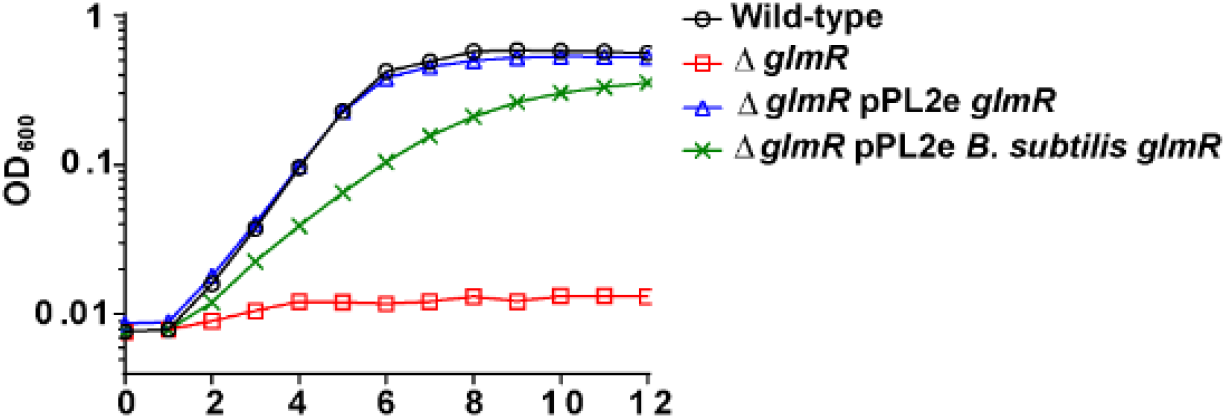
*B. subtilis* GlmR rescues the cell wall defect of a Δ*glmR* mutant. Transcomplementation of growth in BHI with 1mg/mL lysozyme over 12 hours at 37°C. Graph is representative of greater than 3 biological replicates.

**Figure S7.**
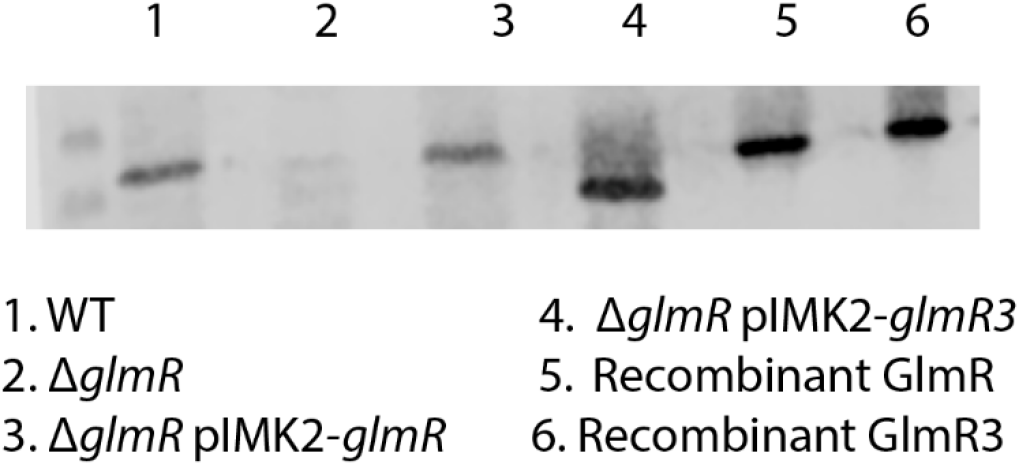
GlmR3 equal or increased expression to WT GlmR. Expression of GlmR in WT, Δ*glmR*, Δ*glmR:glmR*, and Δ*glmR:glmR3* at mid-log in BHI with 250 ug/mL lysozyme.

**Table S1.**
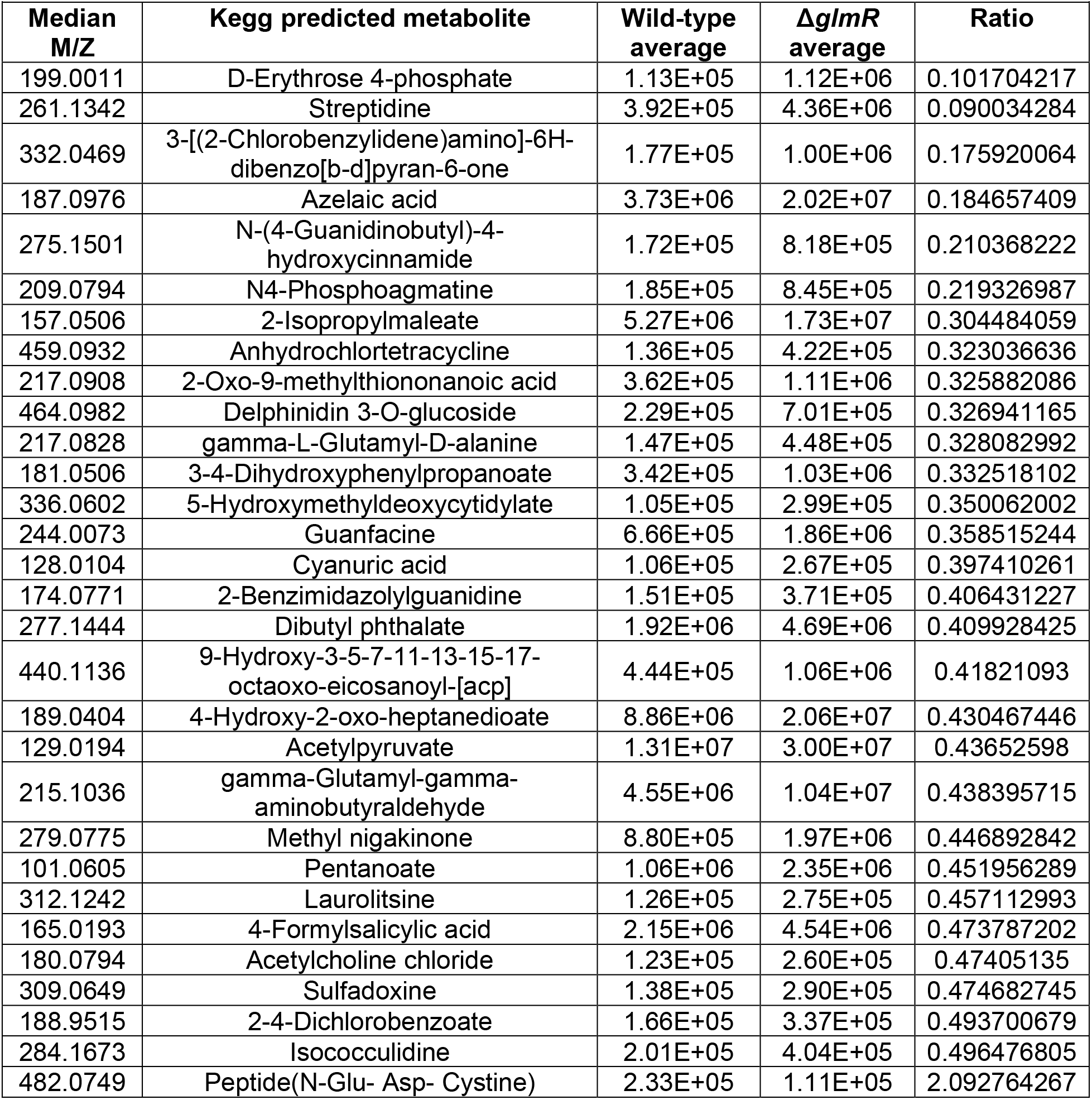
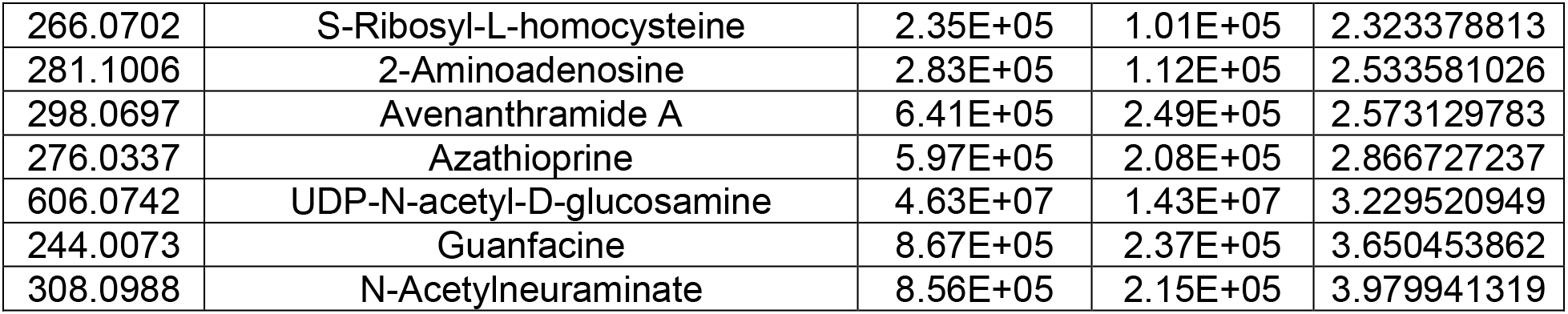
Putative KEGG identified differential metabolites. Putative KEGG identified metabolites with greater than 2-fold abundance differential between wild-type and Δ*glmR*, their m/z, abundance in wild-type and the *glmR* mutant, and ratio between the two are listed.

## References

1. Goetz M, Bubert A, Wang G, Chico-Calero I, Vazquez-Boland JA, Beck M, Slaghuis J, Szalay AA, Goebel W. 2001. Microinjection and growth of bacteria in the cytosol of mammalian host cells. Proc Natl Acad Sci U S A 2001/09/27. 98:12221–12226.

2. Goetz M, Engelbrecht F, Goebel W. 2004. Inefficient Replication of Listeria innocua in the Cytosol of Mammalian Cells. J Infect Dis 393–401.

3. Brumell JH, Rosenberger CM, Gotto GT, Marcus SL, Finlay BB. 2001. SifA permits survival and replication of Salmonella typhimurium in murine macrophages. Cell Microbiol 3:75–84.

4. Chen GY, McDougal CE, D’Antonio MA, Portman JL, Sauer J-D. 2017. A Genetic Screen Reveals that Synthesis of 1,4-Dihydroxy-2-Naphthoate (DHNA), but Not Full-Length Menaquinone, Is Required for Listeria monocytogenes Cytosolic Survival. MBio 8:e00119–17.

5. Creasey EA, Isberg RR. 2012. The protein SdhA maintains the integrity of the Legionella-containing vacuole. Proc Natl Acad Sci 109:3481–3486.

6. Sampson TR, Napier BA, Schroeder MR, Louwen R, Zhao J, Chin C-Y, Ratner HK, Llewellyn AC, Jones CL, Laroui H, Merlin D, Zhou P, Endtz HP, Weiss DS. 2014. A CRISPR-Cas system enhances envelope integrity mediating antibiotic resistance and inflammasome evasion. Proc Natl Acad Sci 111:11163–11168.

7. Peng K, Broz P, Jones J, Joubert LM, Monack D. 2011. Elevated AIM2-mediated pyroptosis triggered by hypercytotoxic Francisella mutant strains is attributed to increased intracellular bacteriolysis. Cell Microbiol 13:1586–1600.

8. Portnoy DA, Chen* C, Mitchell* G. 2016. Strategies Used by Bacteria to Grow in Macrophages. Microbiol Spectr 4:1–22.

9. Mitchell G, Isberg RR. 2017. Innate Immunity to Intracellular Pathogens: Balancing Microbial Elimination and Inflammation. Cell Host Microbe 22:166–175.

10. Casanova JE. 2017. Bacterial Autophagy: Offense and Defense at the Host–Pathogen Interface. Cell Mol Gastroenterol Hepatol 4:237–243.

11. Eisenreich W, Dandekar T, Heesemann J, Goebel W. 2010. Carbon metabolism of intracellular bacterial pathogens and possible links to virulence. Nat Rev Microbiol 8:401–12.

12. Liu W, Zhou Y, Peng T, Zhou P, Ding X, Li Z, Zhong H, Xu Y, Chen S, Hang HC, Shao F. 2018. Nε-fatty acylation of multiple membrane-associated proteins by Shigella IcsB effector to modulate host function. Nat Microbiol 3:996–1009.

13. Cheng MI, Chen C, Engström P, Portnoy DA, Mitchell G. 2018. Actin-based motility allows *Listeria monocytogenes* to avoid autophagy in the macrophage cytosol. Cell Microbiol 20:e12854.

14. Sauer J-D, Witte CE, Zemansky J, Hanson B, Lauer P, Portnoy D a. 2010. Listeria monocytogenes triggers AIM2-mediated pyroptosis upon infrequent bacteriolysis in the macrophage cytosol. Cell Host Microbe 7:412–9.

15. Reniere ML, Whiteley AT, Portnoy DA. 2016. An In Vivo Selection Identifies Listeria monocytogenes Genes Required to Sense the Intracellular Environment and Activate Virulence Factor Expression. PLOS Pathog 12:e1005741.

16. Pensinger DA, Boldon KM, Chen GY, Vincent WJB, Sherman K, Xiong M, Schaenzer AJ, Forster ER, Coers J, Striker R, Sauer J-D. 2016. The Listeria monocytogenes PASTA Kinase PrkA and Its Substrate YvcK Are Required for Cell Wall Homeostasis, Metabolism, and Virulence. PLOS Pathog 12:e1006001.

17. Smith HB, Li TL, Liao MK, Chen GY, Guo Z, Sauer J-D. 2021. Listeria monocytogenes MenI encodes a DHNA-CoA thioesterase necessary for menaquinone biosynthesis, cytosolic survival, and virulence. Infect Immun 89:e00792–20.

18. Görke B, Foulquier E, Galinier A. 2005. YvcK of Bacillus subtilis is required for a normal cell shape and for growth on Krebs cycle intermediates and substrates of the pentose phosphate pathway. Microbiology 151:3777–91.

19. Mir M, Prisic S, Kang C-M, Lun S, Guo H, Murry JP, Rubin EJ, Husson RN. 2014. Mycobacterial gene cuvA is required for optimal nutrient utilization and virulence. Infect Immun 82:4104–17.

20. Griffin JE, Gawronski JD, DeJesus M a., Ioerger TR, Akerley BJ, Sassetti CM. 2011. High-resolution phenotypic profiling defines genes essential for mycobacterial growth and cholesterol catabolism. PLoS Pathog 7:1–9.

21. Chaudhuri RR, Allen AG, Owen PJ, Shalom G, Stone K, Harrison M, Burgis TA, Lockyer M, Garcia-Lara J, Foster SJ, Pleasance SJ, Peters SE, Maskell DJ, Charles IG. 2009. Comprehensive identification of essential Staphylococcus aureus genes using Transposon-Mediated Differential Hybridisation (TMDH). BMC Genomics 10:291.

22. Foulquier E, Galinier A. 2017. YvcK, a protein required for cell wall integrity and optimal carbon source utilization, binds uridine diphosphate-sugars. Sci Rep 7:4139.

23. Barreteau H, Kovač A, Boniface A, Sova M, Gobec S, Blanot D. 2008. Cytoplasmic steps of peptidoglycan biosynthesis. FEMS Microbiol Rev 32:168–207.

24. Brown S, Santa Maria JP, Walker S. 2013. Wall Teichoic Acids of Gram-Positive Bacteria. Annu Rev Microbiol 67:313–336.

25. Jankute M, Grover S, Birch HL, Besra GS. 2014. Genetics of Mycobacterial Arabinogalactan and Lipoarabinomannan Assembly. Microbiol Spectr 2.

26. Patel V, Wu Q, Chandrangsu P, Helmann JD. 2018. A metabolic checkpoint protein GlmR is important for diverting carbon into peptidoglycan biosynthesis in Bacillus subtilis. PLOS Genet 14:e1007689.

27. Promadej N, Fiedler F, Cossart P, Dramsi S, Kathariou S. 1999. Cell wall teichoic acid glycosylation in Listeria monocytogenes serotype 4b requires gtcA, a novel, serogroup-specific gene. J Bacteriol 181:418–25.

28. Eugster MR, Morax LS, Hüls VJ, Huwiler SG, Leclercq A, Lecuit M, Loessner MJ. 2015. Bacteriophage predation promotes serovar diversification in Listeria monocytogenes. Mol Microbiol 97:33–46.

29. Foulquier E, Pompeo F, Byrne D, Fierobe HP, Galinier A. 2020. Uridine diphosphate N-acetylglucosamine orchestrates the interaction of GlmR with either YvcJ or GlmS in Bacillus subtilis. Sci Rep 10.

30. Battesti A, Bouveret E. 2012. The bacterial two-hybrid system based on adenylate cyclase reconstitution in Escherichia coli. Methods 58:325–334.

31. Pompeo F, Bourne Y, van Heijenoort J, Fassy F, Mengin-Lecreulx D. 2001. Dissection of the Bifunctional *Escherichia coli N*-Acetylglucosamine-1-phosphate Uridyltransferase Enzyme into Autonomously Functional Domains and Evidence That Trimerization Is Absolutely Required for Glucosamine-1-phosphate Acetyltransferase Acti. J Biol Chem 276:3833–3839.

32. Graupner M, Xu H, White RH. 2002. Characterization of the 2-phospho-L-lactate transferase enzyme involved in coenzyme F(420) biosynthesis in Methanococcus jannaschii. Biochemistry 41:3754–61.

33. Blake KL, O’Neill AJ, Mengin-Lecreulx D, Henderson PJF, Bostock JM, Dunsmore CJ, Simmons KJ, Fishwick CWG, Leeds JA, Chopra I. 2009. The nature of *Staphylococcus aureus* MurA and MurZ and approaches for detection of peptidoglycan biosynthesis inhibitors. Mol Microbiol 72:335–343.

34. Foulquier E, Pompeo F, Bernadac A, Espinosa L, Galinier A. 2011. The YvcK protein is required for morphogenesis via localization of PBP1 under gluconeogenic growth conditions in Bacillus subtilis. Mol Microbiol 80:309–18.

35. Pouliot Y, Karp PD. 2007. A survey of orphan enzyme activities. BMC Bioinformatics 8:244.

36. Sévin DC, Fuhrer T, Zamboni N, Sauer U. 2016. Nontargeted in vitro metabolomics for high-throughput identification of novel enzymes in Escherichia coli. Nat Methods.

37. de Carvalho LPS, Zhao H, Dickinson CE, Arango NM, Lima CD, Fischer SM, Ouerfelli O, Nathan C, Rhee KY. 2010. Activity-based metabolomic profiling of enzymatic function: identification of Rv1248c as a mycobacterial 2-hydroxy-3-oxoadipate synthase. Chem Biol 17:323–32.

38. Rae CS, Geissler A, Adamson PC, Portnoy DA. 2011. Mutations of the Listeria monocytogenes peptidoglycan N-deacetylase and O-acetylase result in enhanced lysozyme sensitivity, bacteriolysis, and hyperinduction of innate immune pathways. Infect Immun 2011/07/20. 79:3596–3606.

39. Sharma R, Khan IA. 2016. Mechanism and potential inhibitors of GlmU: a novel target for antimicrobial drug discovery. Curr Drug Targets.

40. Foulquier E, Pompeo F, Freton C, Cordier B, Grangeasse C, Galinier A. 2014. PrkC-mediated phosphorylation of overexpressed YvcK protein regulates PBP1 protein localization in Bacillus subtilis mreB mutant cells. J Biol Chem 289:23662–9.

41. Mignolet J, Viollier PH. 2011. A sweet twist gets Bacillus into shape. Mol Microbiol 80:283–285.

42. Monk IR, Gahan CGM, Hill C. 2008. Tools for functional postgenomic analysis of listeria monocytogenes. Appl Environ Microbiol 74:3921–3934.

43. Pensinger DA, Aliota MT, Schaenzer AJ, Boldon KM, Ansari IH, Vincent WJB, Knight B, Reniere ML, Striker R, Sauer J-D. 2014. Selective pharmacologic inhibition of a PASTA kinase increases Listeria monocytogenes susceptibility to β-lactam antibiotics. Antimicrob Agents Chemother 58:4486–94.

44. Zemansky J, Kline BC, Woodward JJ, Leber JH, Marquis H, Portnoy DA. 2009. Development of a mariner-based transposon and identification of Listeria monocytogenes determinants, including the peptidyl-prolyl isomerase PrsA2, that contribute to its hemolytic phenotype. J Bacteriol 191:3950–64.

45. Burke TP, Loukitcheva A, Zemansky J, Wheeler R, Boneca IG, Portnoy D a. 2014. Listeria monocytogenes Is Resistant to Lysozyme through the Regulation, Not the Acquisition, of Cell Wall-Modifying Enzymes. J Bacteriol 196:3756–3767.

46. Hodgson DA. 2000. Generalized transduction of serotype 1/2 and serotype 4b strains of Listeria monocytogenes. Mol Microbiol 2000/01/29. 35:312–323.

47. Sun AN, Camilli A, Portnoy DA. 1990. Isolation of Listeria monocytogenes small-plaque mutants defective for intracellular growth and cell-to-cell spread. Infect Immun 58:3770–3778.

48. Luu P, Gorman T. 1997. Short communication A chemically defined minimal medium of Listeria for the optimal culture. Int J Food Microbiol 35:91–95.

49. Whiteley AT, Garelis NE, Peterson BN, Choi PH, Tong L, Woodward JJ, Portnoy DA. 2017. c-di-AMP modulates Listeria monocytogenes central metabolism to regulate growth, antibiotic resistance and osmoregulation. Mol Microbiol 104:212–233.

50. Rydzak T, Garcia D, Stevenson DM, Sladek M, Klingeman DM, Holwerda EK, Amador-Noguez D, Brown SD, Guss AM. 2017. Deletion of Type I glutamine synthetase deregulates nitrogen metabolism and increases ethanol production in Clostridium thermocellum. Metab Eng 41:182–191.

51. Clasquin MF, Melamud E, Rabinowitz JD. 2012. LC-MS data processing with MAVEN: a metabolomic analysis and visualization engine. Curr Protoc Bioinformatics Chapter 14:Unit14.11.

52. Melamud E, Vastag L, Rabinowitz JD. 2010. Metabolomic Analysis and Visualization Engine for LC−MS Data. Anal Chem 82:9818–9826.

53. Sun L, Rogiers G, Michiels CW. 2021. The Natural Antimicrobial *trans*-Cinnamaldehyde Interferes with UDP-N-Acetylglucosamine Biosynthesis and Cell Wall Homeostasis in *Listeria monocytogenes*. Foods 10, 1666.

